# MTBP allosterically activates Cdk8-CycC kinase activity

**DOI:** 10.1101/2025.06.16.659917

**Authors:** Eman Zaffar, Julio Vieyto-Nuñez, Katerina Pravi, Mohammed Faidh Aslam, Yasser Almeida-Hernández, Joel Mieres-Pérez, Farnusch Kaschani, Markus Kaiser, Anika Kroesen, Dominik Boos, Elsa Sánchez-García

## Abstract

How the approximately 300 human protein kinases identify their dedicated substrates despite the significant overlap of their phosphorylation consensus sequences is relevant for nearly all cellular processes. We show here that the Cdk8/19-CycC kinase uses mutually exclusive targeting and activation factors to facilitate distinct cellular roles. The Med12 protein is known to target Cdk8/19-CycC to the mediator of transcription complex to control the transcription of specific gene sets upon respective stimuli. We describe that a second Cdk8/19-CycC targeting factor exists, the replication origin firing regulator MTBP that targets the kinase to Med12-independent cellular roles. Both Med12 and MTBP constitute allosteric activators of the enzymatic Cdk8/19-CycC kinase activity in vitro. We describe the structural basis of this activation that involves distinct mechanisms how Med12 and MTBP reposition the T-loop of the kinase independently of T-loop phosphorylation – the canonical mechanism of CDK kinase activation. Our results support the following model: the Cdk8/19-CycC dimer alone has low enzymatic activity, which may help avoid off-target phosphorylation. Med12, MTBP and potentially other as yet unidentified accessory factors, target the kinase to distinct molecular environments, at the same time activating kinase activity for efficient substrate phosphorylation. Our work establishes an unusual mechanism of CDK kinase control and change the current paradigm how Cdk8/19-CycC selects its substrates. Substrate selection of the kinase may be relevant for cancer biology and therapy because Cdk8 is a well-established colorectal cancer promoting factor.

## Introduction

Virtually all cellular processes are controlled by kinases that regulate their protein substrates by phosphorylation of serine, threonine or tyrosine amino acids ^1^. The roughly 300 kinases of the human proteome must phosphorylate exactly their right substrate sites and only in the right conditions. The eukaryotic serine/threonine kinases are part of only three large kinase families, basophilic, acidophilic and proline-directed kinases. Structural similarities in their catalytic sites underlie this classification, mediating preference for basic or acids acidic in the vicinity of the phospho-residue, or for proline following the phospho-residue. The structural similarities create a significant overlap of intrinsic enzymatic substrate specificities between the kinases within a family, although their specificities are typically not completely identical. This specificity overlap necessitates other mechanisms to direct kinases to their designated substrate sites. Among such mechanisms, specific docking to substrate proteins through short linear sequence motifs (SLIMs) in the kinase substrate is common ^2^. For a docking mechanism to work effectively, the kinase should possess limited phosphorylation capacity before docking - be it through low kinase protein levels or low catalytic activity - to prevent significant background phosphorylation and to ensure that only the docked substrate is modified.

Cdk8/19-cyclin C (CycC) is a member of a branch of Cyclin-dependent kinases (CDKs) CDK kinases that have roles in gene transcription (tCDKs). CDK are proline-directed kinases comprising a heterodimer of one kinase and one regulatory cyclin subunits ^3^. 20 CDK kinases are known in the human proteome (CDK1-CDK20), grouped into three groups based on their function in the cell. The first group (CDK 1, 2, 3, 4, and 6) is mostly involved in the regulation of the cell cycle, the second group are tCDKs (CDK 7, 8, 9, 10, 11, 12 and 13). In the third group are CDKs with atypical features and functions (CDK 5, and 14 to 20), like non-canonical activation mechanisms for example by non-cyclin proteins and membrane association ^4–8^. Full activity of canonical CDKs requires association with their cyclin subunits and phosphorylation of the T-loop in the kinase subunit, allowing the entry of the substrate into the catalytic site.^3,9^

tCDKs control gene transcription at several levels. Cdk8 associates with CycC to control transcriptional activation or repression ^10^. Cdk8-CycC is widely conserved in eukaryotes. In mammals, Cdk8 has a close relative, Cdk19, that has overlapping but distinct roles with Cdk8 ^11^. Cdk8-CycC is involved in the tumorigenesis and progression of colorectal cancer, melanoma, breast and prostate cancer, as well as uterine leiomyoma ^12–18^. Cdk8/19-CycC associates with the Med12 and Med13 proteins to form the CDK kinase module (CKM) of the mediator of transcription complex that regulates the recruitment of transcription and regulation factors that modify the activity of Pol II, in a context-dependent manner.^19^ Whereas mediator controls transcription of almost all PolII-transcribed genes, Cdk8/19-CycC regulates specific sets of genes upon stimulation ^10^. Examples are the transcriptional response to stimulation by interferon gamma (IFNγ), where Cdk8/19-CycC phosphorylates STAT1 at serine 727 to induce STAT1- dependent gene expression ^11,20^ and the control of glycolytic genes in hypoxic conditions ^21^. The enzymatic kinase activity of Cdk8/19-CycC is required for many cellular roles of the kinase, although kinase independent roles have been suggested ^10,22,23^ ^11,21^. The relevant kinase substrates have been identified in only few cases. In complex with Med12, Cdk8/19-CycC phosphorylates the C-terminal domain of RNA polymerase II (RNA-PolII-CTD). RNA-PolII-CTD comprises dozens of repeats of the consensus sequence YSPTPS (or variants thereof): The different serines, threonines and tyrosines of the repeats are subject to differential phosphorylation by several kinases, the pattern of which on the long CTD constitutes a phosphorylation code to control the RNA-PolII transcription cycle ^24^.

Canonical CDKs contain a conserved threonine residue (T160 in CDK2) in the T-loop whose modification is required for kinase activity ^25^. Hence these kinases are called phosphorylation-dependent CDKs. Phosphorylation repositions the T-loop, creating a network of interactions based on three conserved arginines that stabilize the T-loop, allowing the substrate peptide to enter the binding site. The segment RHYTK, which is situated a few amino acids C-terminal of the T-loop, also participates in the stabilization of the T-loop of Cdk2, with tyrosine residue Y180 (Y211 in Cdk8) being particularly important ^26^. Cdk8 lacks the T-loop threonine residue, making this phosphorylation unnecessary for the full activation of Cdk8-CycC. T-loop repositioning is instead mediated by interaction with Med12^27^. Residues in the N-terminal 100 amino acids of Med12 interact and stabilize with the T-loop and the RHYTK segment ^26,28,29^, allowing the substrate to enter the catalytic site.

We recently identified the MTBP protein as an interactor of Cdk8/19-CycC ^30^. MTBP forms a regulatory hub for replication origin firing together with its complex partners Treslin and TopBP1 ^31^. This complex mediates cell cycle regulation of replication, coupling replication origin firing to the S phase ^32,33^. Moreover, the MTBP-Treslin-TopBP1 helps control origin firing in replication stress conditions that occur upon DNA damage and oncogene activation by inhibiting origin firing to avoid mutation through ^32–37^. MTBP and its regulation is therefore important to maintain genetic stability in normal and adverse growth conditions. In line with this regulatory role in replication, we found that the interaction between MTBP and Cdk8/19-CycC is required to prevent replication stress in cultured human cells ^30^. The molecular and cellular mechanisms how MTBP-Cdk8/19-CycC exerts this stress protection function remain unclear. We speculate that the interaction controls replication origin firing in space, time and/or frequency to support replication fidelity.

Here, we use computational structural analysis and biochemical re-constitution to show that MTBP is an allosteric activator of the enzymatic activity of Cdk8-CycC. We suggest a molecular mechanism of this activation that resembles that of activation by Med12, but that also shows mechanistic distinctions. Our work contributes molecular insight of the role of MTBP-Cdk8/19-CycC, which constitutes an important step to understand the role of the complex in replication. Our identification of MTBP as a Cdk8/19-CycC activator also changes the paradigm how Cdk8/19-CycC acts in cells. Our model suggests that the kinase uses alternative targeting and activation factors - MTBP, Med12 and potentially others - for distinct cellular roles.

## Results and discussion

### Mutually exclusive binding of MTBP and Med12 to Cdk8-CycC

We described that MTBP-containing complexes of Cdk8-CycC immunoprecipitated from cell lysates did not contain Med12 and Med13 ^30^. The mutually exclusive binding may arise through direct competition of MTBP and Med12/Med13 interaction surfaces on the kinase. To test if the Cdk8/19-CycC binding domains of MTBP ^26,30^ and Med12 ^26,28^ were sufficient for the mutual competition, we over-expressed either binding domain at increasing levels in Hek293T cells. We co-overexpressed 6Myc-Cdk8 for subsequent immunoprecipitation by anti-Myc antibodies. The immunoprecipitation enriched for Myc-Cdk8 as well as for endogenous full-length MTBP and Med12 (Fig 1A, Fig S1). Overexpression of the N-terminal 100 amino acids of Med12 (Med12(N100)) reduced the amount of co-precipitated endogenous Med12 and MTBP. The same was true when MTBP-516-704, from which the Treslin binding N-terminal domain and the C-terminal Sld7-like domain had been deleted, was overexpressed. In contrast, a mutant of MTBP-516-704 that cannot bind Cdk8/19-CycC binding due to seven amino acid exchanges (MTBP-mut for binding mutant) did not compete with the endogenous MTBP and Med12 proteins ^30^. This result is consistent with direct competition between MTBP and Med12 for

**Figure 1:**
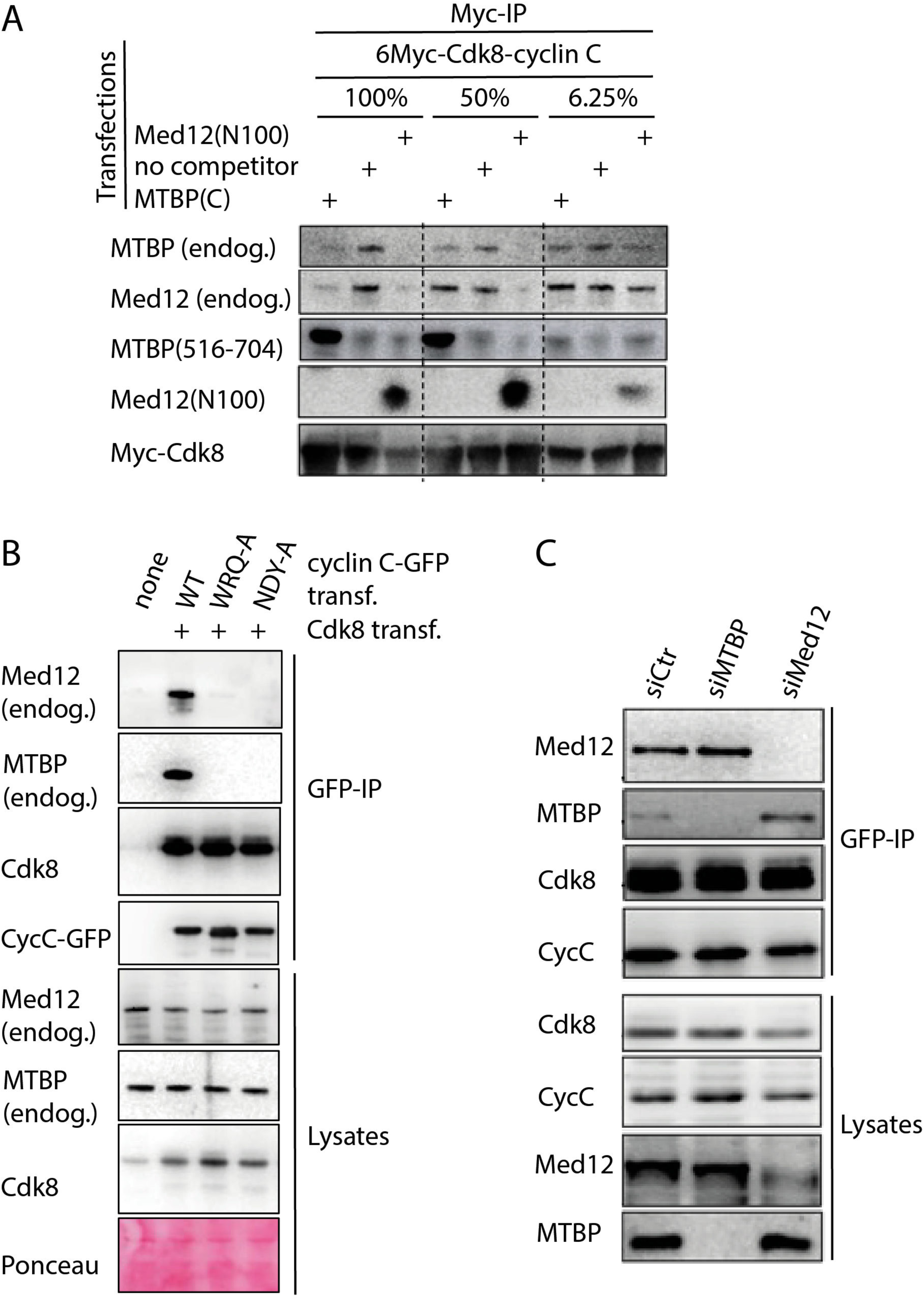
Mutually exclusive binding of MTBP and Med12 to Cdk8-cyclin. **C** A)Co-immunoprecipitation experiment with 6Myc-Cdk8 from lysates of Hek293T cells transfected with wild-type (WT) or mutant 6Myc-Cdk8 and Cdk8-cyclin C binding fragments of Med12 (Med12(N100)), MTBP (MTBP-516-704-WT) or the non-Cdk8-cyclin C binding mutant of MTBP (MTBP-516-704-mut). B) GFP-co-immunoprecipitation of CycC-3Flag-GFP-WT or the indicated point mutants upon transient overexpression in Hek293T cells. Cdk8 was co-overexpressed. C) Anti-Cdk8 immunoprecipitation from Hela cell lysates upon treatment with siRNAs against MTBP (siMTBP), Med12 (siMed12) or with control siRNA (siCtr).

Cdk8-CycC binding. Next, we tested if MTBP and Med12 use the same binding surfaces on the Cdk8/19-CycC kinase. Analysis of recently published structures of the yeast and human Cdk8-CycC kinase bound to Med12 suggested that human CycC residue triplet W6, R58 and Q59 as well as triplet N181, D182 and Y238 form important interactions with Med12(N100) residues. We mutated these triplets separately to alanine and tested binding to endogenous Med12 and MTBP. For this, C-terminally 3Flag-GFP-tagged CycC-WT or the mutants (MTBP-WRQ-A, NDY-A) were transfected into Hek293T cells followed by anti-GFP immunoprecipitation. Figure 1B shows that Cdk8, MTBP and Med12 coprecipitated with CycC-WT. In contrast, both CycC mutants showed strongly reduced interactions with both Med12 and MTBP, whereas they interacted normally with Cdk8. This experiment suggests that Med12 and MTBP use at least partly the same binding surfaces on CycC, consistent with a direct competition mechanism.

We wanted to know next if the level of endogenous MTBP in cells affects the level Cdk8/19-CycC kinase binding to Med12 and vice versa. We used siRNAs to deplete MTBP and Med12, respectively, from Hela cells and asked if the knock-downs changed the amount of the non-depleted protein bound to the kinase. Anti-Cdk8 immunoprecipitation from cell lysates showed that siMed12 prevented co-immunoprecipitation of Med12 with Cdk8 and siMTBP co-immunoprecipitation of MTBP (Fig 1C). siMed12 led to slightly increased levels of MTBP bound to the kinase, and siMTBP increased kinase-bound Med12. Albeit weak, these effects were reproducible. Together, the co-immunoprecipitation studies presented in Figures 1A-C suggested that the MTBP and Med12 are in a competitive relationship regarding binding to the Cdk8-CycC kinase in cultured human cells.

We then tested whether established cellular functions of Cdk8/19-CycC depended on Med12 or MTBP, respectively. The mutually exclusive binding observed led us to expect dependency on Med12 or MTBP, but not on both simultaneously. We first tested the interferon gamma (IFNγ) response, which is known to be partly dependent on Cdk8/19-CycC phosphorylation of STAT1-S727 upon treatment with IFNγ ^11,20^. siRNAs against Cdk8, CycC and Med12 decreased STAT1-S727 phosphorylation in response to IFNγ treatment in western blots of whole cell lysates (Fig 2A). In contrast, siMTBP did not have an effect. This is consistent with the scenario that, for IFNγ−induced STAT1 phosphorylation, Cdk8-CycC acts as part of the mediator kinase module with Med12 and Med13, and that MTBP is dispensable. We then tested the expression of the glucose transporter SLC2A3. Our qPCR analysis confirmed earlier observations that SLC2A3 expression was induced by hypoxic conditions and that treatment with the Cdk8/19 inhibitor (Cdk8i) MSC2530818 reduced expression in hypoxic and also in normorxic growth conditions in HCT116 cells (Fig 2B) ^21^. This shows dependency of SLC2A3 expression on Cdk8-19-CycC enzymatic activity. Consistently, siRNA treatment against Cdk8 reduced transcript levels of SLC2A3 and Cdk8 (Fig 2C). Surprisingly, MTBP siRNA reduced SLC2A3 mRNA (i), suggesting that MTBP is required for normal SLC2A3 expression. MTBP siRNA did not change Cdk8 mRNA levels, showing that the effect of siMTBP on SLC2A3 was not caused by decreased Cdk8 (ii). In contrast siMTBP, siMed12 treatment increased SLC2A3 mRNA levels. This could be due to increased levels of MTBP bound to Cdk8/19-CycC (Fig 1C) upon depletion of Med12 protein, which could induce SLC2A3 expression. Thus, expression of SLC2A3 is independent of Med12 and requires MTBP for high expression levels in HCT116 cells.

**Figure 2:**
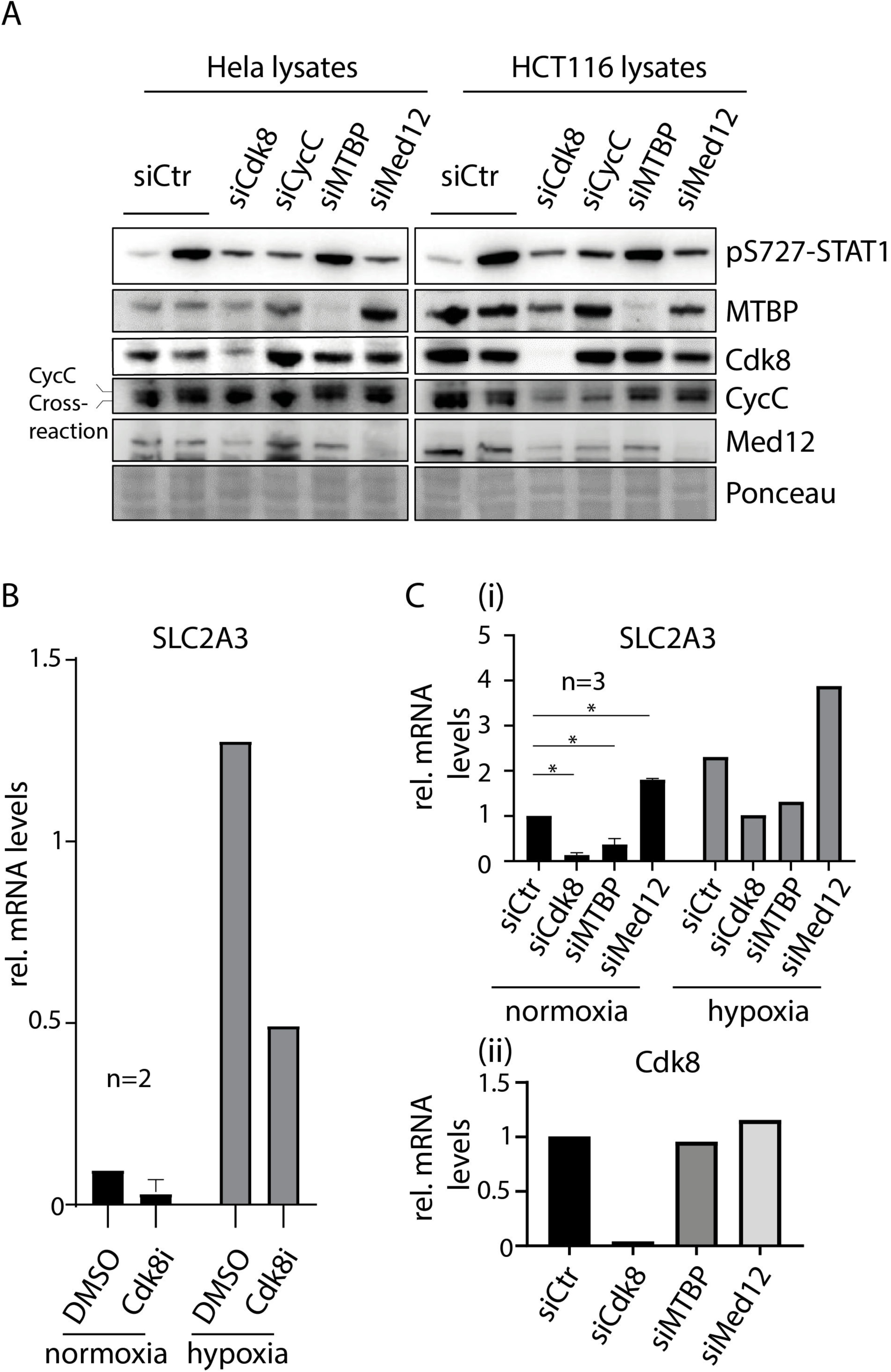
Distinct roles of Cdk8-CycC together with Med12 and MTBP. A) Interferon gamma (IFNγ) induced STAT1-pS727 phosphorylation depends on Cdk8-CycC and Med12, but not on MTBP. IFNγ-treated and control-treated cells were lysed for immunoblotting with the indicated antibodies. B) SLC2A3 mRNA levels are controlled by Cdk8 kinase activity in normoxic and hypoxic conditions. HCT116 cells were treated with the Cdk8 inhibitor MSC2530818 before mRNA extraction and quantitative RT-PCR. C) The control of SLC2A3 mRNA levels does not require Med12, but Cdk8 and MTBP, as assessed by qRT-PCR. Treatment with control siRNA (siCtr) or siRNAs against Cdk8, MTBP or Med12 was done for 72h before mRNA extraction and qRT-PCR to analyze the levels of SLC2A3 (i) and Cdk8 (ii).

Together, these experiments clearly separate Cdk8/19-CycC functions in cells in Med12-dependent and independent functions and support mutual competition of Med12 and MTBP in cells for Cdk8/19-CycC interaction.

### Interaction of MTBP and Med12 with CycC

We performed extensive microsecond-scale GaMD simulations to gain insight into the conformational dynamics of CycC when complexed with MTBP or Med12. The binding of both MTBP and Med12 to CycC results in stabilization of large regions of CycC (residues 50-60, 100-126, 155-175, and 190-230) with respect to the unbound Cdk8-CycC complex, as evident from differences in the residue-level root mean square fluctuation (RMSF) profiles (Fig 3A). The RMSF values in the regions from amino acids 50-60 and 100-126 of CycC are affected to a similar extent by both MTBP and Med12. In contrast, MTBP affects the CycC regions formed by residues 155-175 and residues 190-230 to a larger extent than Med12 (Fig 3A). Those differences can be rationalized based on distinct interaction paths of both Cdk8-CycC binders along CycC (Fig. 3B). Furthermore, one segment of MTBP binds to an area of CycC located at the border of the regions formed by residues 155-175 and 190-230 of CycC, and interacts with residues in both regions simultaneously. In contrast, Med12 interacts with residues of those regions separately (Fig. 3B). This could explain why the regions of CycC comprising residues 155-175 and 190-230, are more rigid when CycC is bound to MTBP than when bound to Med12.

**Figure 3:**
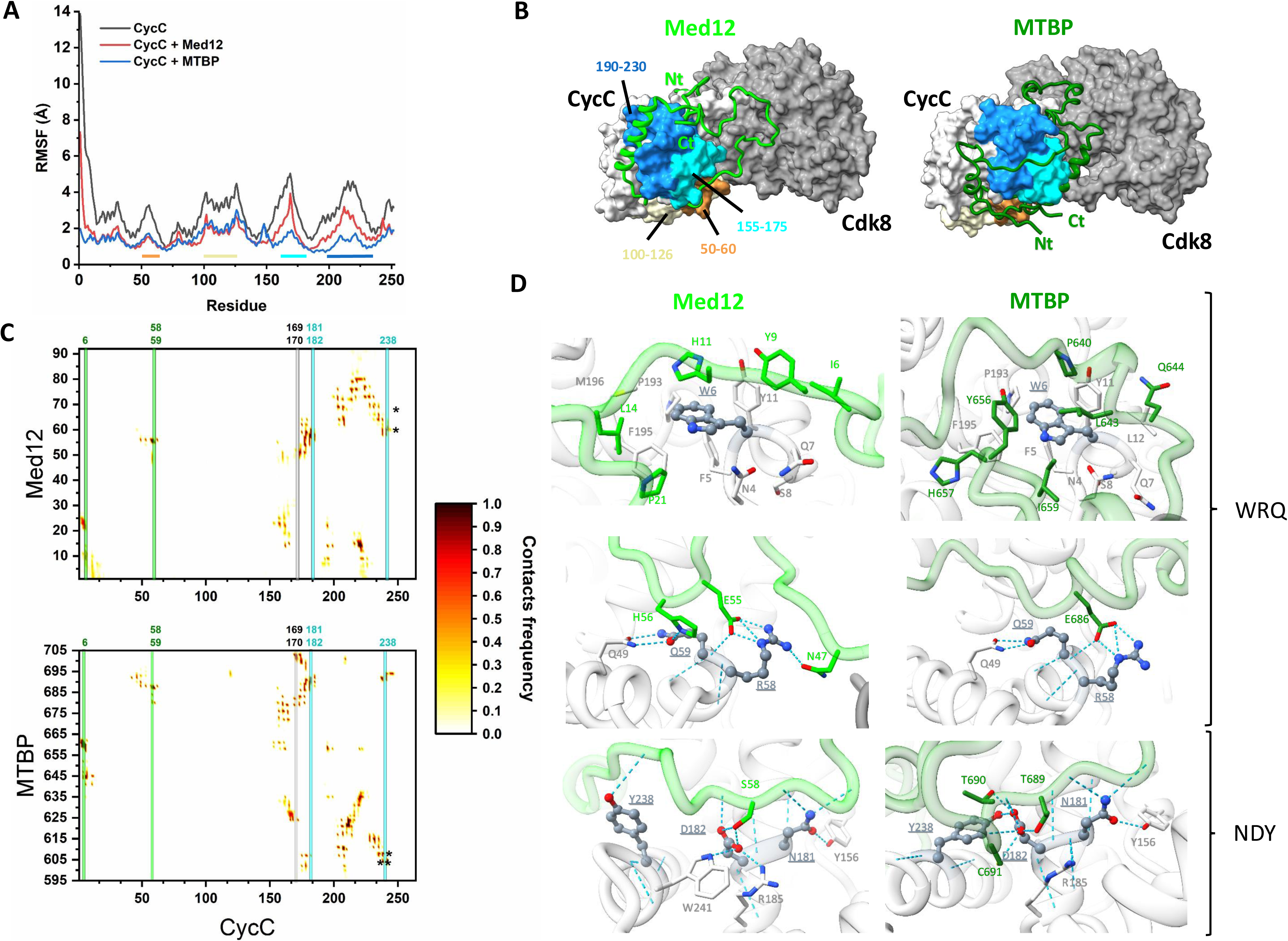
Similar, but distinct, interaction of Med12 and MTBP with CycC. A) Residue-level root mean square fluctuation (RMSF) profiles of CycC, bound to Med12 (red line) and MTBP (blue line). The coloured bars below the curves highlight different segments of residues of CycC that show changes upon MTBP and/or CycC binding. Orange: 50-60, pale yellow: 100-126, cyan: 155-175, blue: 190-230. B) Representative structure of Cdk8 (light gray)-CycC (dark gray) complex, bound to Med12(N100) (light green) and MTBP(595-705) (dark green). The colored surface regions correspond to the coloured segments highlighted in panel A. C) 2D contact maps between residues of CycC and Med12(N100) (top) and MTBP(595-705) (bottom). The color scale represents the contact frequency of the heavy atoms of the residues (cutoff distance = 5 Å), during the simulations. The black asterisks mark residue pairs identified to form BS3-crosslinks (Fig S2). D) Neighboring residues of the CycC-WDR-A and CycC-NFY-A triple mutants. The mutated residues are represented in ball and sticks. The hydrogen bonds and salt bridges interactions are represented as cyan, dashed lines.

To verify this computational insight, we used BS3-mediated crosslinking-mass spectrometry. This analysis confirmed the region 190-230 of CycC that contains residues with high interaction frequencies both with Med12 and MTBP (Fig S2 and Fig 3C, black asterisks). For MTBP, but not Med12, also a crosslink in the computationally predicted interaction region aa 50-60 region could be detected (K56), whereas the other predicted interaction regions in CycC did not result in crosslinks. For further validation of the GaMD simulation results, we mutated residues predicted to be important for the interaction of MTBP and Med12 with Cdk8-CycC. For interaction studies the mutated CycC were transiently expressed in Hek293T cells followed by for GFP-immunoprecipitation from lysates as described for Figure 1B. As discussed above, mutation to alanines of the combinations WRQ (W6, R5, Q59) and NDY (N181, D282, Y238) of CycC strongly affected the binding of both MTBP and Med12 (Fig 1B). This is reflected in the high contact frequency of these residues with both kinase binders throughout the simulations, and by the nature of the interactions involved (Fig 3C,D). In the CycC-WRQ-A triple mutant, the W6A mutation affects a hydrophobic patch formed by surrounding CycC residues (F5, Y11, L12, P193, F195) with Med12 residues I6, Y9, L14, P21, and MTBP residues P640, L643, Y56, and I659, respectively (Fig 3D, top panels). The CycC-R58A mutation affects two main interactions with Med12, a conserved salt bridge between residues R58 and E55 (contact frequency = 97%) and a transient hydrogen bond between residues R58 and N47 (contact frequency = 47%) (Fig 3D, middle panel). This mutation also affects a salt bridge between the same arginine residue from CycC and and residue E686 from MTBP (contact frequency = 100%) (Fig 3D, middle panel). Furthermore, CycC residue Q59 establishes hydrogen bond interactions with residue CycC-Q49 and Med12-H56 (Fig 3D). Hence, the hydrophobic patch of CycC-W6 and the salt bridges of CycC-R58 constitute key interactions of CycC with Med12 and MTBP, respectively. In the CycC-NDY-A triple mutant, the mutated N181, D182 and Y238 residues of CycC are close to CycC-R58/Q59 and establish several hydrogen bonds with Med12 residues 55-62 and MTBP residues 686-691 (contact frequencies > 95%) (Fig 3D, bottom panels). These results reveal the presence of two main anchor points for the interaction of MTBP and Med12 with CycC: the hydrophobic patch around CycC-W6 and the hydrophilic patch comprising CycC-R58, Q59, N181, D182, Y238.

We then mutated residues that should selectively affect the binding of CycC to Med12 but not MTBP. Indeed, the selected quadruple mutant CycC-YFDM (Y192A, F217A, D223A, M224A) affected the binding to Med12 stronger than the binding to MTBP (Fig 4A). These CycC residues belong to the region 190-230 mentioned above (Fig 3A,B), and form a hydrophobic cleft together with several other CycC residues (L119, L202, A213, A218, L220, V222, I227, L228, I231, V233, I234, L235, L237) (Fig. 4B). This hydrophobic cleft accommodates an α-helix containing residues Med12-L14, F73, I76 and I77 with a predicted contact frequency larger than 80% during the GaMD simulations (Fig 3C, Fig 4B). Additionally, residue D223 from CycC establishes two transient salt bridges with residues from Med12. One salt-bridge with arginine R12 (contact frequency = 47%) and the other one with lysine K80 (contact frequency = 28%). MTBP on the other hand, does not interact with those residues during the GaMD simulations, which could explain the lower impact of the Y192A, F217A, D223A, M224A mutations on the binding of CycC to MTBP compared to that of Med12. Together, the mutation analysis presented support the computational predictions of the MTBP-Cdk8-CycC complex.

**Figure 4:**
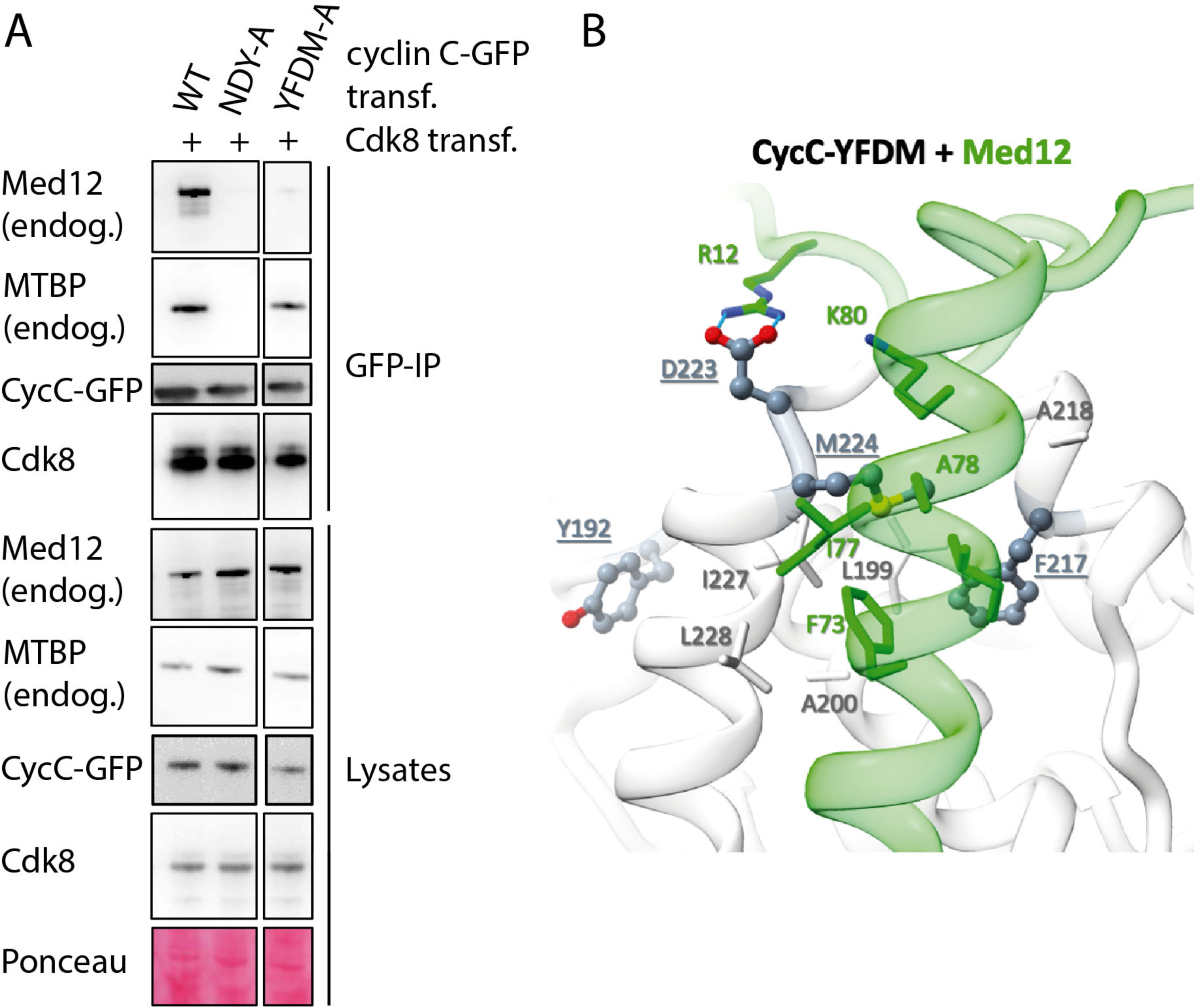
Distinct effects of the CycC-YDFM mutant on MTBP and Med12 binding are consistent with computational structural predictions. A) Co-immunoprecipitation experiment from Hek293T cell lysates as described in Figure 1B using the indicated CycC mutants. B) Structural model of the interactions between Med12(N100) and the interaction cleft in CycC containing Y192A, F217A, D223A and M224A that are mutated in the CycC-YFDM mutant.

These results show that MTBP and Med12 use overlapping, but distinct, interaction surfaces on CycC. Med12(N100) and MTBP(C) bear little sequence and structural similarity, if any, which is highlighted by the opposite N-terminal to C-terminal orientation with which the two peptides run along with the CycC surface (Fig 3B).

### Distinct modes of Cdk8 t-loop stabilization by MTBP and Med12

Next, we analyzed the interactions of Med12 and MTBP with the Cdk8 subunit. GaMD simulations suggested that Cdk8 exhibits pronounced stabilization in several regions upon binding of MTBP and Med12, including the T-loop region (residues 180–187) and the RHYTK segment (residues 209–213) (Fig 5A), which could also be recapitulated by BS3-mediated crosslinks (Fig S2). Both T-loop and RHYTK motif, constitute critical regulatory regions for Cdk8 kinase activity. The RHYTK motif plays a pivotal role in stabilizing the T-loop into an active conformation ^26^. Structural studies have demonstrated that Med12 binds to the RHYTK segment, facilitating T-loop stabilization through specific contacts, thereby promoting kinase activation ^26^. Our GaMD simulations show that binding of both MTBP and Med12 lead to stabilization of Cdk8 residues 108–126, 175–195 (including the T-loop). The T-loop undergoes a significant reduction in mobility upon binding, with RMSF values falling below 6 Å for the holo forms (Fig 5A). The RHYTK motif displays significant stabilization upon binding of Med12, in contrast to a small stabilizing effect observed with MTBP (Fig 5A). The Cdk8 region comprising residues 240-250 is also more stabilized by Med12 than by MTBP. The 240-250 region is a Cdk8/19-specific insertion between helices αF and αG of the kinase protein that is not found in other Cdk kinases ^38^. These observations establish several interaction regions in Cdk8 for MTBP and Med12, including the T-loop and RHYTK segment. The modes with which MTBP and Med12 interact with these Cdk8 regions show differences that could have functional consequences.

**Figure 5:**
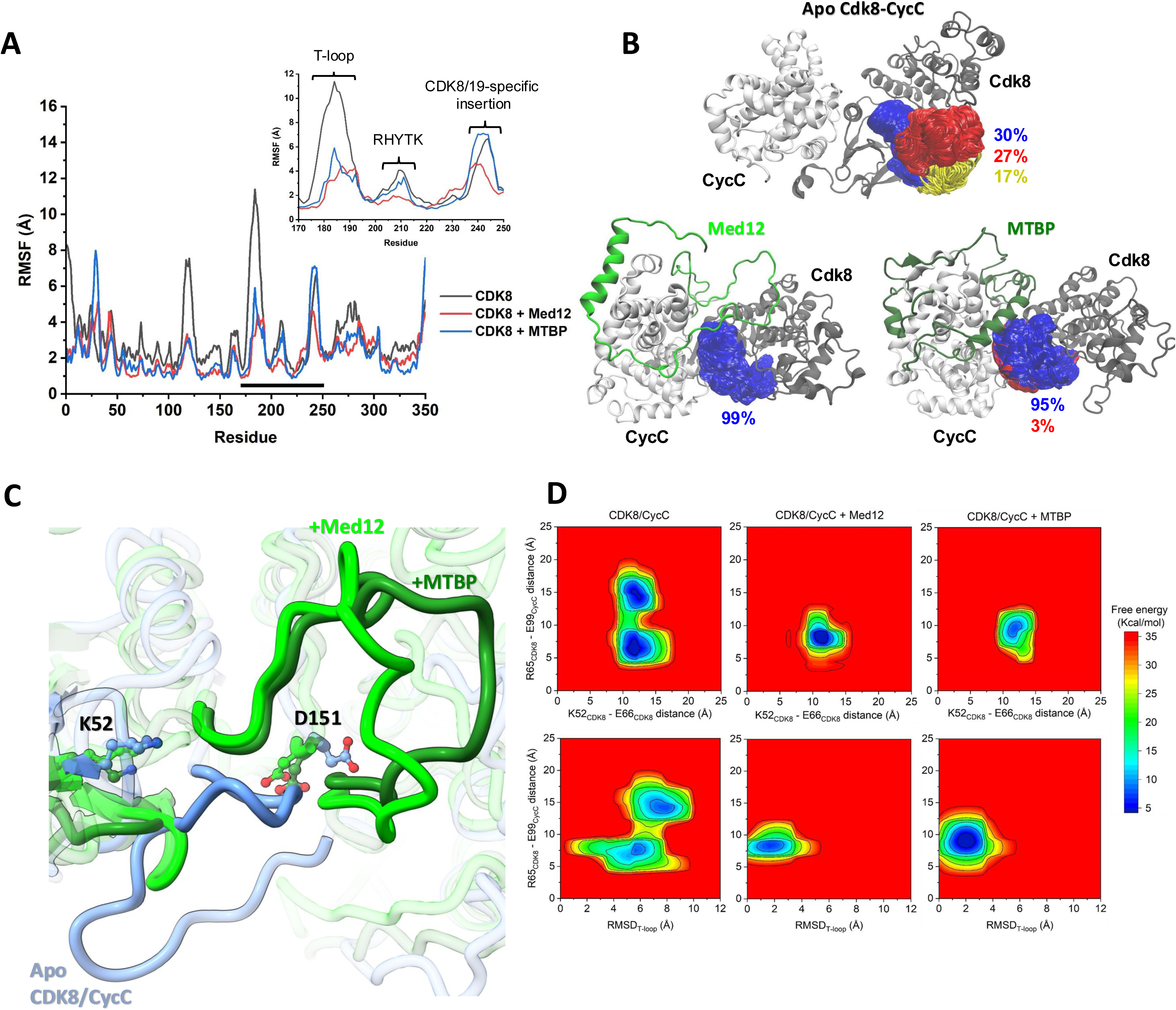
Distinct modes of Cdk8 T-loop stabilization by MTBP and Med12. A) Residue-level root mean square fluctuation (RMSF) profiles of CDK8, bound to Med12 (red line) and MTBP (blue line). The black bar below the curves highlights the segment corresponding to the T-loop, RHYTK segment, and the Cdk8/19-specific insertion (inner plot). B) Clustering analysis of the T-loop in the apo Cdk8-CycC dimer complex, and when bound to Med12 or MTBP. The conformations of the T-loop were clustered with an RMSD cutoff value of 5 Å. The percent values correspond to the size of the cluster with respect to the whole simulation trajectory. C) Conformations of the T-loop in the apo Cdk8-CycC complex (blue), and bound to Med12 (light green) and MTBP (dark green). Representative conformations based on GaMD simulations are shown. Residues K52 and D151 of Cdk8 are represented in licorice. D) Reweighted potential-mean force (PMF) maps of the T-loop conformational sampling, described with different collective variables. Top: Distance between CDK8-K52@Cα and CDK8-E66@Cα vs distance between in CDK8-R65@Cα and CycC-E99@Cα. Bottom: RMSD of the T-loop‘s backbone vs. distance between CDK8-R65@Cα and CycC-E99@Cα.

The effects of the Cdk8-CycC binders MTBP and Med12 on T-loop mobility were corroborated by clustering analysis of the T-loop conformations sampled during the GaMD simulations. The apo state of Cdk8-CycC, the kinase dimer with neither Med12 nor MTBP bound, displayed a broad conformational distribution, with the three most populated clusters representing 30%, 27%, and 17% of the trajectory frames, respectively (Fig 5B). Presence of MTBP or Med12 led to a striking reduction in conformational variability: the most populated cluster encompasses 95% and 99% of the explored conformations respectively (Fig 5B), confirming that both MTBP and Med12 restrict the conformational flexibility of the T-loop. In addition, when neither MTBP nor Med12 are bound, the active site of the kinase is occluded by the T-loop, blocking substrate binding. Upon binding of Med12 or MTBP, this occlusion of the active site is removed by a conformational change of the T-loop, allowing the binding of the substrate (Fig 5C). Thus, despite the observed differences in T-loop and RHYTK motif interactions MTBP and Med12 have similar effects on the active kinase site.

To further assess how Med12 and MTBP binding modulates the conformational landscape of the T-loop of Cdk8, we computed various potential mean-force maps (PMFs) along selected collective variables (CVs) (Fig 5D). In the apo form (no MTBP and Med12 bound), the PMF maps display multiple minima, reflecting the large flexibility of the T-loop due to a wide range of accessible conformations. In contrast, both Med12- and MTBP-bound systems show a marked narrowing of the free energy basins, with a single, well-defined minimum dominating the landscape. These results support the model that - despite the observed differences in T-loop and RHYTK motif interactions - both MTBP and Med12 stabilize the active conformation of Cdk8 by restricting T-loop flexibility in a conformation that favors the binding of the substrate. These findings also support the scenario that, like Med12, MTBP acts as an allosteric activator of Cdk8-CycC.

Furthermore, the simulations reveal that Med12 is more flexible than MTBP when bound to Cdk8-CycC, especially in the region comprised by Med12 residues 24-50 (Fig 6A) allowing Med12 to establish a larger number of contacts with Cdk8 than MTBP (Fig 6B,C). These more numerous contacts of Med12 occur, however, with lower frequency. This is reflected by the top-10 most frequent contacts of MTBP and Med12, respectively. The top-10 most frequent contacts of Med12 range from 63% to 93%, while MTBP contacts range from 98% to 100%. Notably, while both MTBP and Med12 show contacts with the T-loop (residues 180–187, Fig 6B, light red box), Med12 establishes contacts with the RHYTK motif (residues 209–213) more frequently than MTBP (Fig 6B, yellow box). Our simulations show that Med12 interacts with the T-loop and the RHYTK segment via several hydrogen bonds and salt bridges, in agreement with the earlier described activation of the yeast Cdk8-CycC kinase ^26^. Specifically, Med12 can establish salt bridges and hydrogen bonds with T-loop residues K185 and D189 residues and RHYTK residues R209 and H210 (Fig 6D). Furthermore, Med12, but not MTBP, also entertains hydrogen bonds and salt bridges with an additional segment formed by residues 268-276 (segment EDIKKMPEH, Fig 6B, cyan box). These Med12-specific interactions could be confirmed by BS3 crosslinking (Fig S2 and Fig 6B, black asterisks). This segment is located far from the Cdk8-CycC interface where most of the interactions are found, suggesting that the same Med12 residues with high RMSF values (aa 24-50) can form contacts with distant surfaces on Cdk8 (T-loop, RHYTK, EDIKKMPEH), showing the flexibility of Med12. MTBP forms prominent contacts between its residues 662-676 and the Cdk8 T-loop region. Cdk8 residues N181, S182, P183 form hydrogen bonds with MTBP residues D672, G673 and S676 in full-length MTBP (contact frequency > 70%) suggesting that MTBP may stabilize the T-loop through direct persistent interactions. The contacts of MTBP with the RHYTK are rather limited, occurring mostly between K213 of Cdk8, and MTBP residue D664 (contact frequency = 80%) (Fig 6D). Interestingly, MTBP does not interacts with the EDIKKMPEH segment of Cdk8, showing a more rigid interface with CycC than Med12. These findings suggest that, although MTBP and Med12 both stabilize the T-loop region, the stabilization occurs via different mechanisms. Whereas MTBP stabilizes the T-loop mostly by direct interaction, Med12 engages more flexibly with both T-loop and RHYTK segment.

**Figure 6:**
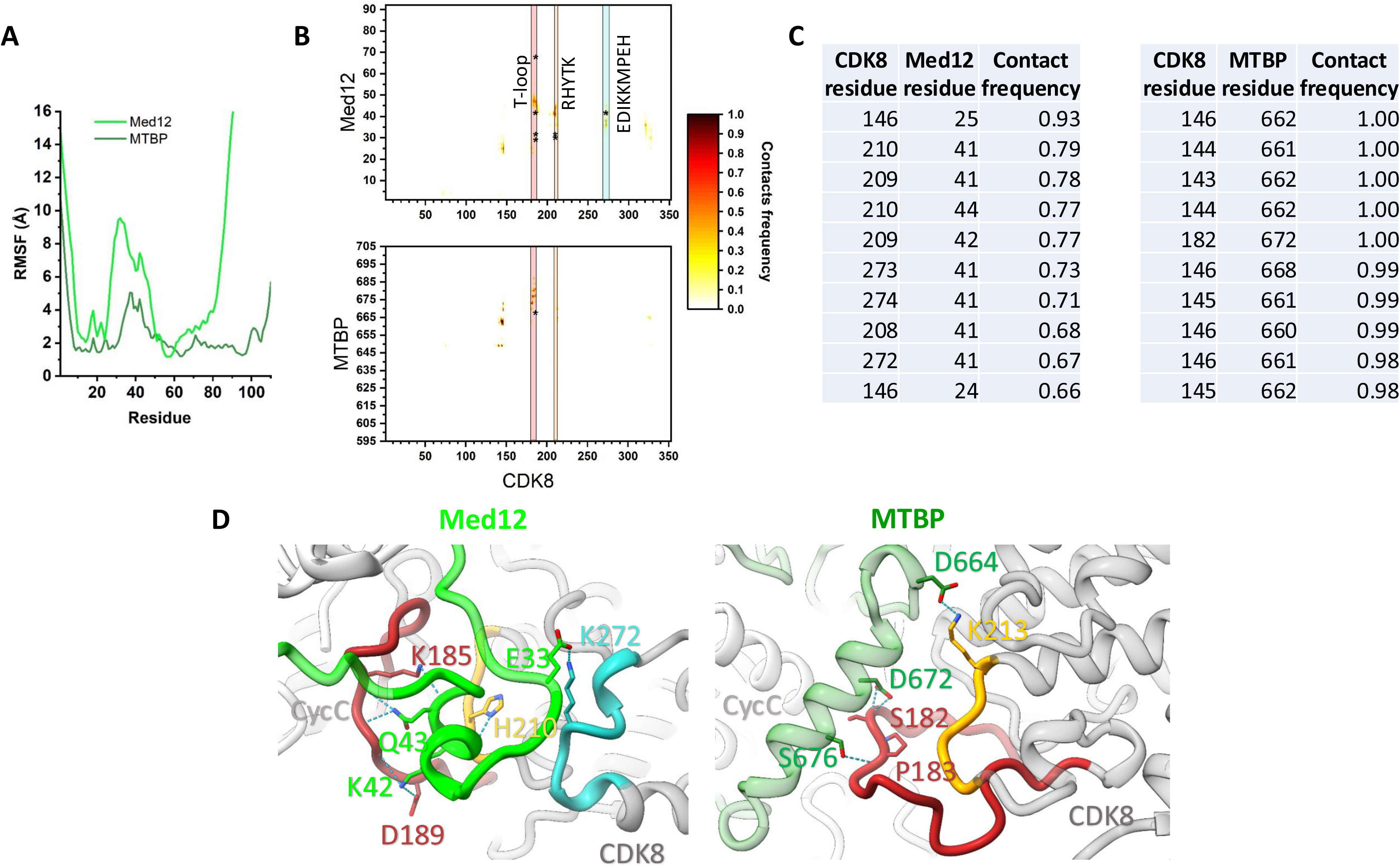
Detailed characterization of Cdk8 contacts with Med12 and MTBP. A) Residue-level root mean square fluctuation (RMSF) profiles of Med12(N100) (light green line) and MTBP-595-704 (dark green line). B) 2D contact map between residues of Cdk8 and Med12 (top) and MTBP (bottom). The color scale represents the contact frequency of the heavy atoms of the residues (cutoff distance = 5 Å), during the simulations. The black asterisks mark residue pairs identified to form BS3-crosslinks (Fig S2). Pink box: T-loop, orange box: RHYTK segment, cyan box: EDIKKMPEH segment. C) Top-10 highest contacting residue pairs of Cdk8-CycC bound to Med12 or MTBP. D) Inter-residue interactions of Med12 residues (light green, left), and MTBP (dark green, right), with residues of the T-loop (red), RHYTK segment (yellow), and EDIKKMPEH segment (cyan). Hydrogen bond and salt bridges interactions are represented as dashed, cyan lines.

The different interaction modes of MTBP and Med12 may have functional consequences in the activation of Cdk8-CycC apart from T-loop stabilization. The interactions also restrict the motion of the kinase and cyclin domains, consistent with the reduced RMSF values and the clustering analysis described above. Taken together, the free energy landscapes provide strong evidence that both ligands reduce the conformational heterogeneity of the T-loop and reinforce a more compact and functionally relevant structural ensemble.

These structural observations establish that Med12 and MTBP both interact with Cdk8 regions that are relevant for the catalytic kinase activity, the T-loop and the RHYTK segment. T-loop stabilization positions it in a conformational cluster that allows substrate binding. T-loop stabilization by both MTBP and Med12 suggests that, despite the mechanistic differences, MTBP may constitute an activator of the Cdk8-CycC kinase activity that activates the kinase in a mutually exclusive manner to Med12.

### MTBP activates the enzymatic activity of Cdk8-CycC

We tested next if MTBP has Cdk8-CycC activating biochemical activity. For this, we employed in vitro kinase phosphorylation experiments. We purified Cdk8-CycC from insect cell lysates (Fig 7A). Incubation with recombinant RNA-PolII-CTD fragment in the presence of ATP resulted in phosphorylation of serine 5 of the CTD heptad repeats (Fig 7B). The phospho-signal was sensitive to treatment with Cdk8/19-CycC inhibitor (not shown) and was not seen with a kinase-dead version of the kinase (Fig 7B). Addition of a recombinant Med12(N100) fragment to Cdk8-CycC at equimolar concentrations strongly increased RNA-PolII-CTD serine 5 phosphorylation (Fig 8A), confirming previous results ^28^. Because we could not purify MTBP at a reasonable quality for similar addition experiments, we co-expressed Cdk8, CycC with either the Cdk8-CycC kinase binding fragment of MTBP (MTBP(C), amino acids 516-C) or Med12(N100). Subsequent purification via the N-terminal 6His-tag on Cdk8 yielded trimeric complexes with roughly stoichiometric amounts of all subunits (Fig 7A(ii), (iii)). Further size exclusion chromatography failed due to the MTBP fragment dissociating from the kinase. In vitro phosphorylation using these recombinant complexes subsequently showed that MTBP clearly activates phosphorylation of RNA-PolII serine 5 compared to the Cdk8-CycC dimer alone, because 100 nM of MTBP-Cdk8-CycC produced similar phospho-signals as 500-800 nM Cdk8-CycC dimer alone (Fig 8B). Med12 appears to be more potent in activating RNA-PolII-CTD serine 5 phosphorylation, because 200 nM MTBP-bound kinase was required to yield similar phosphorylation levels as 50 nM Med12-Cdk8-CycC. Also note the pronounced phospho-gel shifts seen with the MTBP-bound kinase (from 500 nM) and with the Med12-bound kinase (from 50 nM) that could not be observed with the Cdk8-CycC dimer alone (Fig 8B, Coomassie staining). These observation firmly establish allosteric activation of recombinant Cdk8-CycC by binding to MTBP.

**Figure 7:**
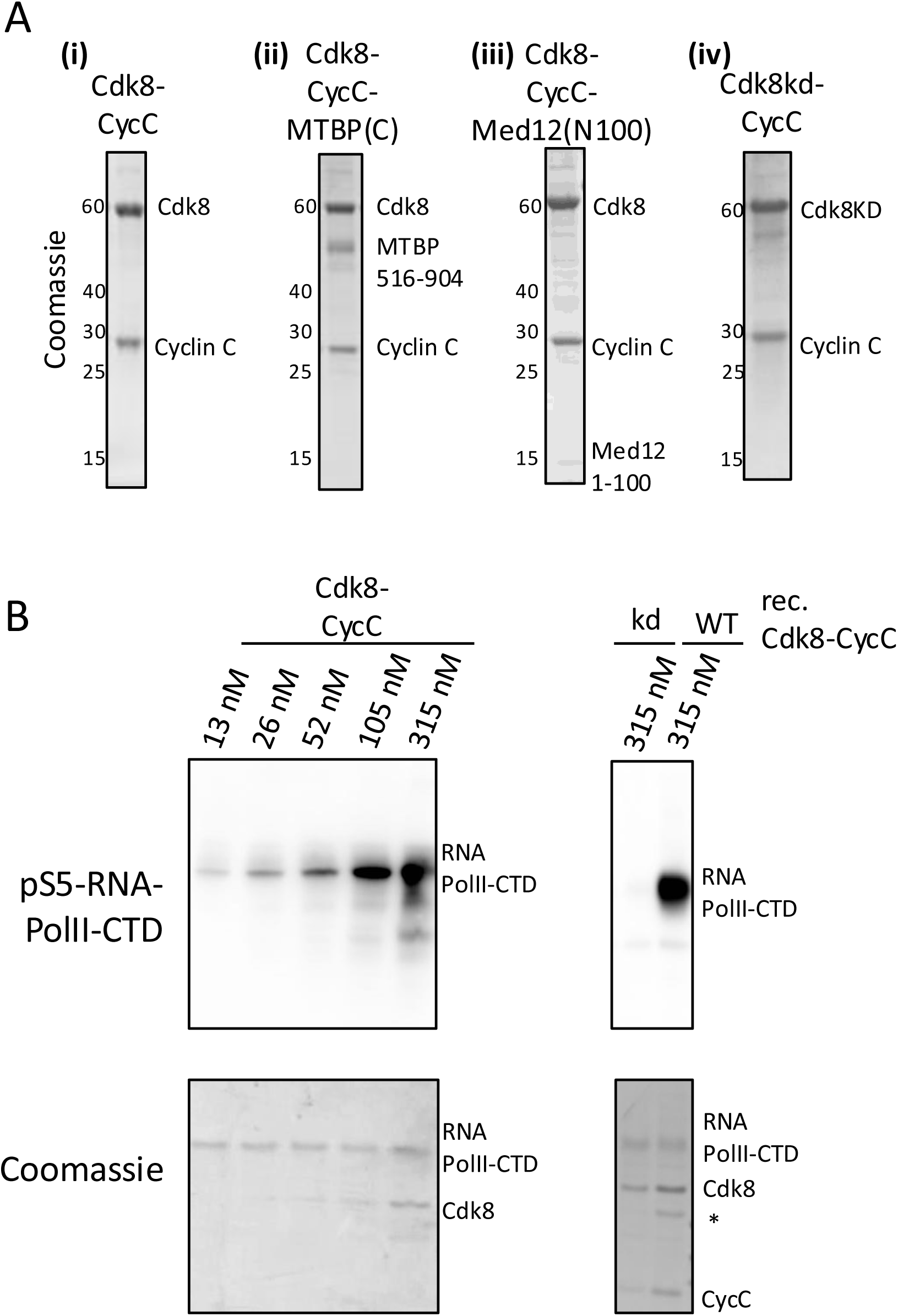
Recombinant Cdk8-cyclin C is an active kinase in vitro. A) Recombinant purified Cdk8-CycC, as a dimer (i) or bound to MTBP-516-C (MTBP(C)) (ii) or to Med12-1-100 (Med12(N100)). The dimeric Cdk8-CycC kinase was also purified as a kinase-inactive (kinase-dead, kd) mutant. B) In vitro phosphorylation experiment using WT or kd Cdk8-CycC versions. RNA-PolI-CTD (C-terminal domain) was used as a substrate. Detection of RNA-PolII-S5 phosphorylation was done by immunoblotting.

**Figure 8:**
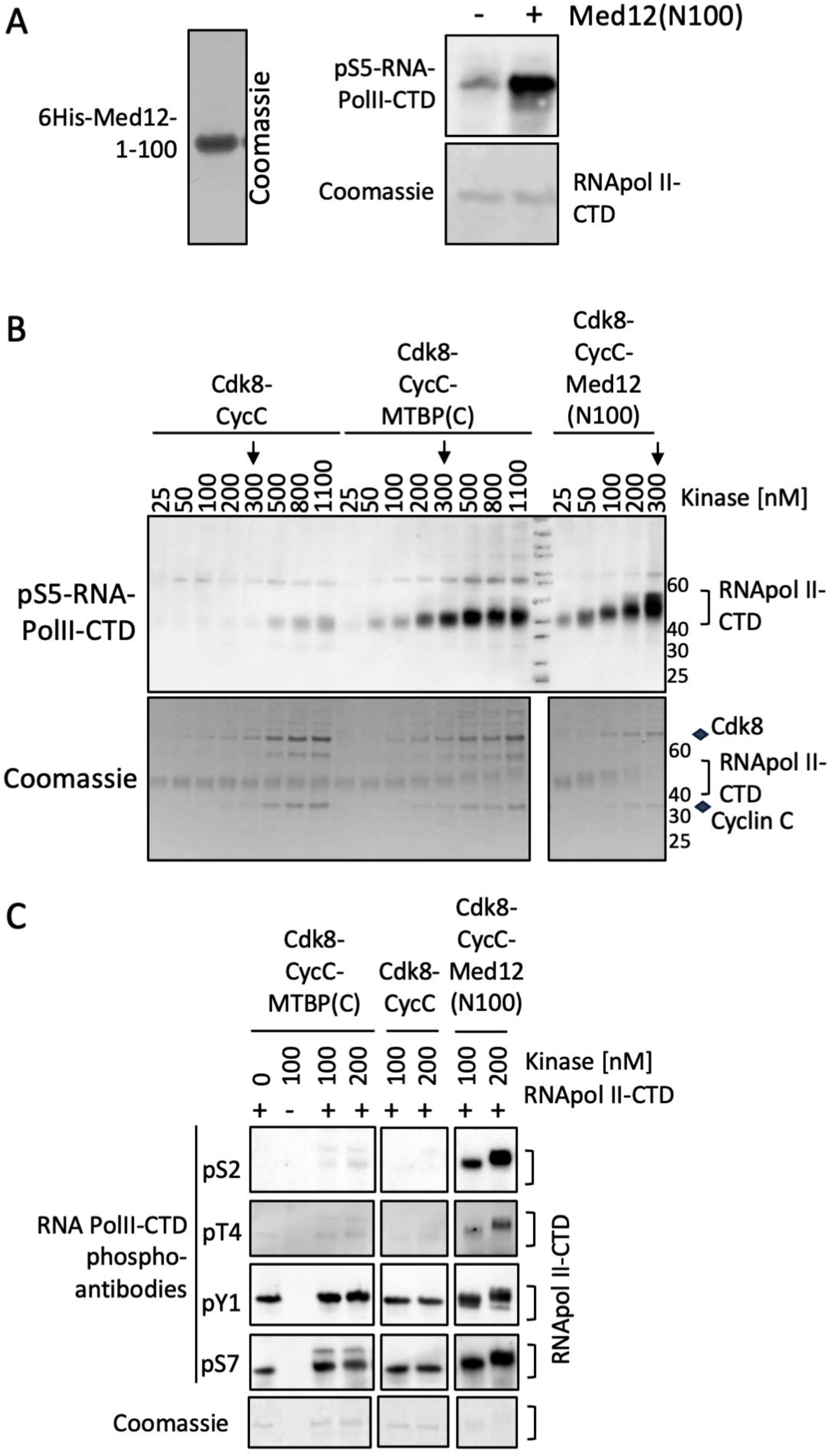
Activation of the enzymatic activity of Cdk8-CycC by MTBP(C) A) In vitro kinase experiment (right panel) as described in Figure 7B in the absence or presence of recombinant Med12(N100) (left panel). B/C) In vitro kinase experiment as described in Figure 7B using recombinant Cdk8-CycC, as a dimer or bound to Med12(N100) or MTBP(C) (Fig 7A). The indicated anti-RNA-PolII-CTD phopsho-specific antibodies were used for B), and C).

To interrogate the differences in activation of RNA-PolII-CTD-serine 5 phosphorylation observed for MTBP(C) and Med12(N100), we tested next phosphorylation of other CTD phospho-site, by immunoblotting using a range of antibodies against CTD-heptad phospho-residues. This revealed that the Med12-bound, but not by the MTBP-bound kinase, phosphorylated CTD-threonine 4 and serine 2, whereas no or low phosphorylation of serine 7 and tyrosine was detected (Fig 8C). This corroborated that both activators, MTBP and Med12, have different effects on the Cdk8-CycC kinase. Whether these differences result from changed substrate specificity or from different overall kinase activity levels remains open.

Together, we conclude that MTBP and Med12 constitute mutually exclusive allosteric activators of the Cdk8-CycC enzymatic kinase activity.

## Conclusion

We here establish MTBP as an allosteric activator of the enzymatic activity of the Cdk8-CycC kinase and suggest a mechanism for this activation. The interaction is dictated by a large binding area on CycC. MTBP uses similar but not identical binding determinants on CycC for the interaction as the only other known allosteric activator of the kinase, Med12. Despite the similarity of the CycC interaction mode, the interaction domains of Med12 and MTBP do not share many structural features, if any, suggesting convergent molecular evolution. A similar pattern emerges for the interaction modes of MTBP and Med12 with the Cdk8 subunit. Both activators stimulate the enzymatic activity through stabilization of the kinase t-loop, independently of phosphorylation of the t-loop, the canonical kinase activation mechanism in other kinases including CDK kinases ^2^. Stabilization of the T-loop by Med12 and MTBP involves equivalent arginine residues in the Cdk8 subunit to form intra-molecular bonds with t-loop residues like those found in Cdk2-cyclin A that bind to the T160-phosphorylated T-loop. However, the aspartate residue in Cdk8 that replaces T160 of Cdk2 is apparently not involved, suggesting that this aspartate does not act by phospho-mimicry in Cdk8. Despite sharing the same end result - T-loop stabilization - MTBP and Med12 act differently, with MTBP establishing extensive contacts with the T-loop directly, whereas Med12 shows strong interactions with the RHYTK segment situated a few amino acids C-terminal of the t-loop. Our data is consistent with the described Med12-dependent activation mechanism of yeast and mammalian Cdk8-CycC ^26,29^.

Binding of proteins to Cdk8-CycC kinase via the CycC subunit is reminiscent of the docking of Cdk-cyclin kinases to their substrates that was described for G1, S and G2M Cdk-cyclin pairs. Several short linear interaction motifs (SLIMS) in the substrate proteins were identified that interact with specific pockets on the G1, S or G2/M-phase cyclin subunits to recruit these cell cycle kinases to their substrates ^39^. Docking helps the kinase to distinguish its designated substrate sites from off-targets. Sometimes docking can even bypass the requirement of phosphorylation for a CDK consensus sequence ^39^. The mechanism employed by Cdk8/19-CycC seems a variation of this scheme (Fig. 9). Here, docking also occurs via specific interaction surfaces on the cyclin subunit CycC. However, Cdk8-CycC docking does not seem to occur predominantly to the substrate to be phosphorylated (although MTBP might be a substrate of Cdk8-CycC ^34^). Instead, docking occurs to an activator and targeting protein that stimulates kinase activity and integrates the kinase into a specific molecular context, mediator in the case of Med12 and MTBP-Treslin-TopBP1 in the case of MTBP. Whether also the intrinsic phosphorylation specificity of Cdk8-CycC changes upon binding to MTBP or Med12, remains to be determined. The dependency of kinase activity on stimulation by MTBP and Med12 may help limit off-target phosphorylation by Cdk8/19-CycC. The integration into the right molecular environment combined with allosteric activation allows phosphorylation of the right kinase substrates. We speculate that MTBP targets Cdk8-CycC to replication origins to control their firing, as suggested by the involvement of the MTBP-Cdk8-CycC interaction in preventing replication stress ^30^. We find here that MTBP might be involved in the induction of the transcription of glucose metabolism genes. Whether MTBP-dependent kinase activation is involved in Cdk8-CycC’s role in tumorigenesis remains to be investigated^16–18^.

**Figure 9:**
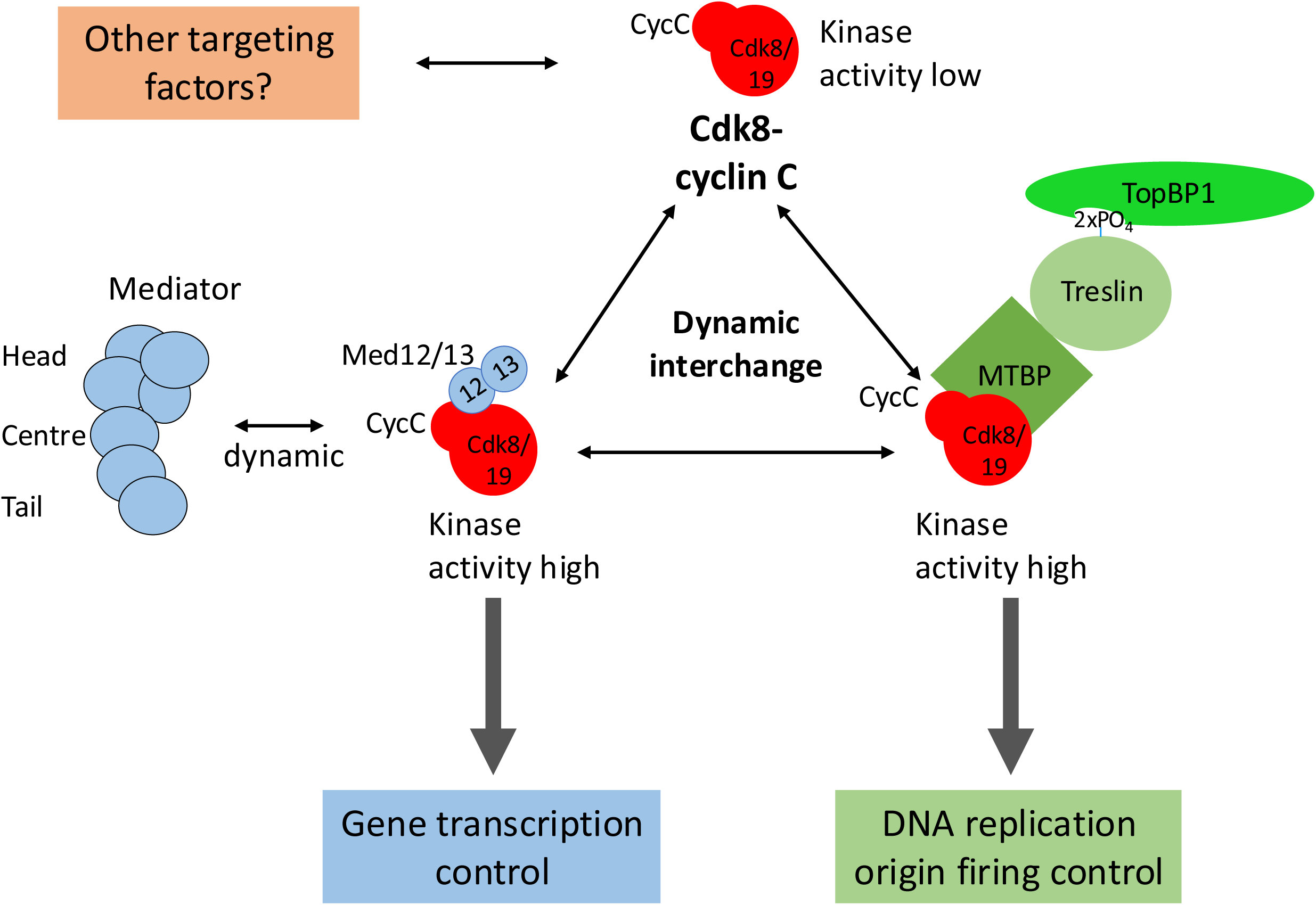
Model for differential targeting of the Cdk8/19-CycC kinase by Med12, MTBP and potentially by other unidentified targeting factors. The Cdk8/19-cyclin C dimer has low enzymatic kinase activity. This may help prevent off-target phosphorylation of CDK consensus sites (S/TP) present in many proteins, for example designated substrates of other proline-directed kinases like cell cycle CDK-regulated proteins or MAP kinase regulated proteins. Mutually exclusive association with Med12 or MTBP activates the Cdk8/19-CycC kinase activity and targets the kinase to distinct molecular environments, to regulate transcription or, presumably, replication origin firing, respectively. Whether other targeting factors exist remains to be investigated.

## Materials and Methods

### Systems preparation

First, we generated a model of human MTBP-Cdk8-CycC and Med12-Cdk8-CycC with AlphaFold2-Multimer ^40^. The models included only the residues 595-704 of MTBP and the N-terminal region of Med12 (residues 1-110). The unfolded, low-scored N-, and C-terminal of Cdk8, CycC, and Med12 were eliminated from the subsequent modeling. As a reference structure for the CDK8/CycC complex, we employed the structure with the PDB ID: 3RGF, ^41^ in which, the missing gaps were added using Modeller.^42,43^ Subsequently, we used CHARMM-GUI web-server^44^ to prepare all the systems using AMBER topology. ^45^The systems were prepared in a cubic simulation box with 10 Å of padding. The box was solvated with water molecules using the TIP3P water model ^46^ ^47^and NaCl at 150 mM was added.

### Gaussian accelerated molecular dynamics (GaMD) simulations

The Cdk8-CycC, MTBP-Cdk8-CycC, and Med12-Cdk8-CycC complexes constructed in the previous step were employed to conduct Gaussian accelerated Molecular Dynamics (GaMD)^48^ in order to enhance the conformational sampling. For Med12, the N-terminal region of residues 1-92 was employed. The CHARMM36m force field^49^ was used and the simulations were carried out in AMBER 24. First, each system underwent energy minimization until the total potential energy dropped below 1000 kcal/mol, followed by NVT equilibration for 500 ps using a 1 fs time step with restraints on the protein’s heavy atoms to allow for solvent relaxation. Next, a 1 ns unrestrained NPT equilibration was performed using 2 fs time step. Subsequently, we carried out the GaMD simulations, beginning with 2.4 ns of cMD and followed by 60 ns of GaMD equilibration to determine boost potential parameters. Finally, three independent dual-boosted GaMD production runs of 1 μs each (3 x 1 μs) were carried out with random initial velocities for the Cdk8-CycC, MTBP-Cdk8-CycC, and Med12-Cdk8-CycC complexes. The simulations were run under periodic boundary conditions, employing the particle mesh Ewald (PME) method for long-range electrostatics. The temperature of 300 K and the pressure of 1 bar were maintained using the Langevin thermostat ^50^ and a Monte Carlo barostat,^51^ respectively.

The trajectories were analyzed using the *cpptraj* tool^39^. Pymol, ChimeraX^52^, and Visual Molecular Dynamics (VMD)^53^ were used for visualization. Conformational clustering of the backbone of the T-loop of CDK8 was performed with the VMD plug-in (https://github.com/luisico/clustering) with a RMSD cutoff value of 5 Å. Contact maps were generated with CONAN with a cutoff distance of heavy atoms of 5 Å ^54^. Energetic reweighting analysis was performed to recover the canonical ensemble from the GaMD simulations using the PyReweighting toolkit ^55^. A set of three collective variables (CVs) was selected to capture key conformational changes associated with the regulatory and catalytic features of the CDK8/CycC complex. These included: (i) the distance between K52 and E66 within CDK8, and (ii) the distance between R65 in CDK8 and E99 in CycC, two critical interactions involved in stabilizing the active conformation of the complex. Additionally, (iii) the RMSD of the activation loop (T-loop) was used to monitor its structural flexibility. The potential of mean force (PMF) was calculated along these CVs by reweighting the biased probability distributions obtained from the GaMD trajectories. The reweighting factor was estimated using a second-order cumulant expansion of the GaMD boost potential, assuming a near-Gaussian distribution of the bias. Finally, two-dimensional PMFs were plotted by combining the CVs into two distinct pairs using Origin Pro ( Version 2023. OriginLab Corporation, https://www.originlab.com).

### Expression construct generation and baculovirus production

Constructs encoding full-length human Cdk8 wt or Kd (N-terminal 6xHis-3xFLAG tagged), human CycC (untagged) were cloned into the pLIB vector. Additional constructs including MTBP wild-type (residues 516–904) with a C-terminal Strep-tag, and human Med12 (residues 1–100), expressed either untagged or with an N-terminal 6xHis-3xFLAG tag. All constructs were confirmed by Sanger sequencing or restriction digestion prior to baculovirus production.

Baculoviruses were generated using FuGENE HD (Promega) mediated transfection of *Spodoptera frugiperda* (Sf9) insect cells. For transfection, approximately 2 × 10^6^ Sf9 cells were seeded per well in a 6-well plate and transfected with 1–2 µg of bacmid DNA mixed with 6 µL of FuGENE HD reagent in serum-free spodopan (PAN BIOTECH). After incubation at 27°C for 3 days, the supernatant containing the P1 virus was collected, clarified by low-speed centrifugation, and stored at 4°C. V1 viral stocks were amplified to produce higher-titer V2 stocks suitable for large-scale infections.

For protein expression, Sf9/Hi5 cells were infected either with individual baculoviruses or co-infected with multiple viruses for complex assembly. Viral infections were performed at a final dilution of 1:50 (virus volume relative to cell culture volume) and the infected cultures were incubated at 27°C with shaking at 100 rpm. Cells were harvested 72 hours post-infection for subsequent protein purification.

### Expression of plasmids in bacterial cells

Constructs encoding human RNA Polymerase II residues 27–52 with an N-terminal 6xHis tag (pJ411 vector, Addgene plasmid #98678) used for bacterial protein expression. DNA constructs were transformed into chemically competent *Escherichia coli* BL21 (DE3) cells according to standard protocols. Following transformation, colonies were selected on LB agar plates containing kanamycin.

For protein expression, a 50 mL LB starter culture supplemented with kanamycin was inoculated with a single colony and grown overnight at 37°C on shaker. The overnight culture was then used to inoculate 1 L of LB medium containing kanamycin at a 1:40 dilution (25 mL starter culture into 1 L LB-kanamycin). Cultures were grown at 37°C until reaching an optical density at 600 nm (OD_600) of 0.6–0.8. Protein expression was induced by the addition of 0.5 mM isopropyl β-D-1-thiogalactopyranoside (IPTG). Following induction, cultures were incubated overnight at 18°C with shaking prior to harvest.

### Purification of His-Tagged Cdk8–Cyclin C, Med12(1–100), RNA pol II 27-52 and Trimeric Kinase Complexes

For purification of His-tagged dimeric Cdk8–Cyclin C wild-type or kinase-dead complexes, Med12(1– 100), and trimeric kinase assemblies, cell pellets were resuspended in lysis buffer composed of 20 mM HEPES (pH 8.0), 150 mM NaCl, 2% (v/v) glycerol, 0.1% (v/v) Triton X-100, 0.5 mM TCEP, 20 mM imidazole, and one protease inhibitor tablet per 50 mL of buffer (Roche Complete, EDTA-free). Cells were lysed by Dounce homogenization (30 strokes) on ice. The lysates were clarified by centrifugation at 20,000g for 30 minutes at 4°C.

The supernatant containing the soluble fraction was incubated with pre-equilibrated Ni-NTA agarose beads for 1 hour at 4°C with gentle rotation. After binding, the beads were transferred to a gravity-flow column and washed three times with wash buffer (identical to the lysis buffer but omitting the protease inhibitors). Bound proteins were eluted using elution buffer containing 20 mM HEPES (pH 8.0), 150 mM NaCl, 2% glycerol, 0.5 mM TCEP, and 250 mM imidazole.

Following elution, proteins were dialyzed overnight at 4°C against dialysis buffer (20 mM HEPES, 100 mM NaCl, 2% glycerol, 0.5 mM TCEP) to reduce salt concentration and remove imidazole.

### In Vitro Kinase Assay

Kinase reactions were carried out in a buffer containing 20 mM HEPES (pH 7.4), 100 mM NaCl, 10 mM MgCl₂, and 2 mM DTT. Substrate proteins were added to the reaction mixture at a final concentration of 100–200 nM. Kinase enzymes were included at concentrations optimized for each experimental condition. Reactions were initiated by the addition ATP to a final concentration of 1 mM and incubated at 30 °C for 10 minutes, or for the durations specified in individual experiments.

To terminate the reactions, an equal volume of 3× Laemmli sample buffer was added, and samples were immediately boiled at 95 °C for 5 minutes. Proteins were resolved by SDS-PAGE and analyzed by western blotting using specific antibodies or by Coomassie Brilliant Blue staining, as appropriate.

### Crosslinking

Crosslinking reactions were performed using BS3 (bis[sulfosuccinimidyl] suberate) in crosslinking buffer composed of 20 mM HEPES (pH 8.0), 150 mM NaCl, 0.01% Tween-20, 2% glycerol, and 0.5 mM DTT. Approximately 10 µM of purified dimeric kinase protein was incubated with either 300 µM or 500 µM BS3, freshly dissolved in the same crosslinking buffer. For trimeric protein complexes, BS3 was used at final concentrations of 750 µM and 1500 µM. Reactions were allowed to proceed at room temperature for 30 minutes. Crosslinking was quenched by the addition of ammonium bicarbonate to a final concentration of 100 mM. Aliquots of the reaction mixtures were analyzed by SDS-PAGE followed by western blotting and silver staining, while the remaining samples were submitted to ACE facilities for crosslinking mass spectrometry (XL-MS).

### RNAi

HeLa Flp-In or HCT116 cells were transfected with siRNAs using RNAiMAX (Life Technologies) according to the manufacturer’s instructions. Briefly, 1.5 × 10⁵ cells per 6-cm tissue culture dish were transfected with 20 nM control siRNA or siRNAs targeting MTBP (1:1 mix of siMTBP-1 and siMTBP-2), CDK8 (1:1 mix of siCDK8-1 and siCDK8-2), Cyclin C (1:1 mix of siCyclin C-1 and siCyclin C-2), or MED12 (1:1 mix of siMED12-1 and siMED12-2), using 10 μL RNAiMAX in a total volume of 5 ml. Cells were harvested 72h post transfection.

### IFN-γ and CDK8 Inhibitors

To assess CDK8 activity in HeLa Flp-In or HCT116 cells, cells were treated with 5 ng/mL IFN-γ for 24 h, resulting in CDK8-mediated phosphorylation of STAT1 at Ser-727. Several CDK8 inhibitors were tested to verify CDK8 specificity: 5 μM Senexin A, 30 μM BRD6989, 2 μM CCT-251921, or 2 μM MSC2530818, for 24 h.

### Cell Culture Conditions for Glycolysis Pathway Experiments

HCT116 cells were treated with control siRNA or siRNAs targeting MTBP, MED12, or CDK8 for 72 h, or with 2 μM MSC2530818 for 24 h. For normoxic conditions, cells were maintained at 37°C in a humidified atmosphere containing 5% CO₂. For hypoxic treatments, cells were incubated at 37°C in a humidified atmosphere containing 1% O₂ and 5% CO₂ for 24 h prior to harvesting.

### qRT-PCR

HCT116 cells were plated and treated as described above followed by harvesting. Total RNA was extracted from cell pellets using RNeasy Plus Mini Kit (Qiagen), according to the manufacturer’s instructions. Reverse transcription was carried out using the Verso cDNA Synthesis Kit (Life Technologies/Thermo Fisher Scientific). Quantitative PCR was carried out with reference to a standard curve using ABsolute qPCR SYBR Green Mix (Life Technologies/Thermo Fisher Scientific) on a CFX96 Touch Real-Time PCR Detection System (BioRad), and normalized to 18S rRNA signals. Primers used were as follows (5’ to 3’): CDK8 fwd – GCAAACAAGAAGCCAGTTCAG, Cdk8 rev – TCTGTGCAACACCCAGTTAG, SLC2A3 fwd – ATGGCCGCTGCTACTGGGTTTT, SLC2A3 rev – ACAACCGCTGGAGGATCTGCTT, 18S fwd – GCCGCTAGAGGTGAAATTCTTG, 18S rev – CTTTCGCTCTGGTCCGTCTT

### Specific materials

#### Plasmids

|  | Company | Reference code |
| --- | --- | --- |
| pGEX-4T-1 3xFlag-STAT1 | Addgene | 82722 |
| RNAPII_CTD27-52 | Addgene | 98678 |

#### Antibodies

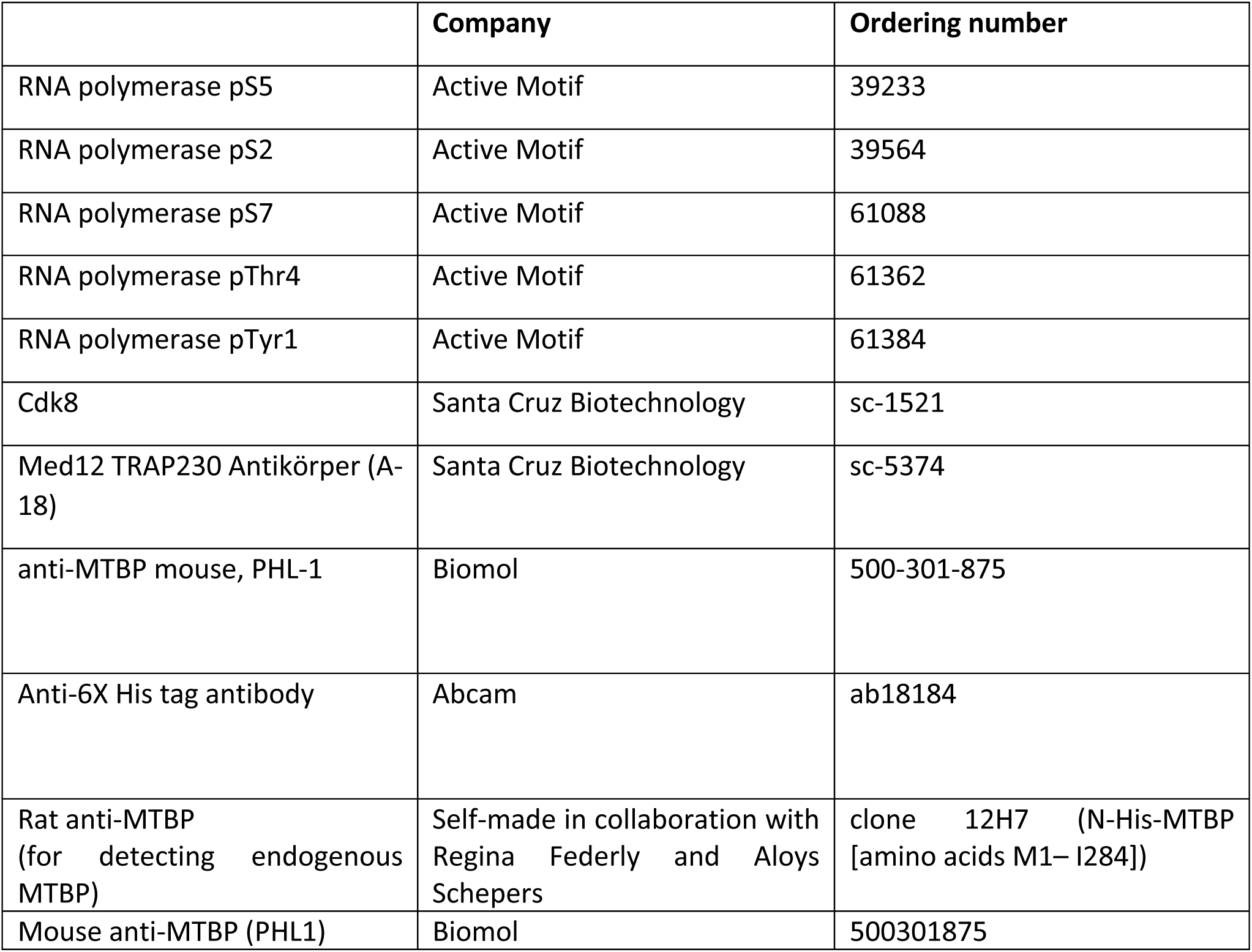
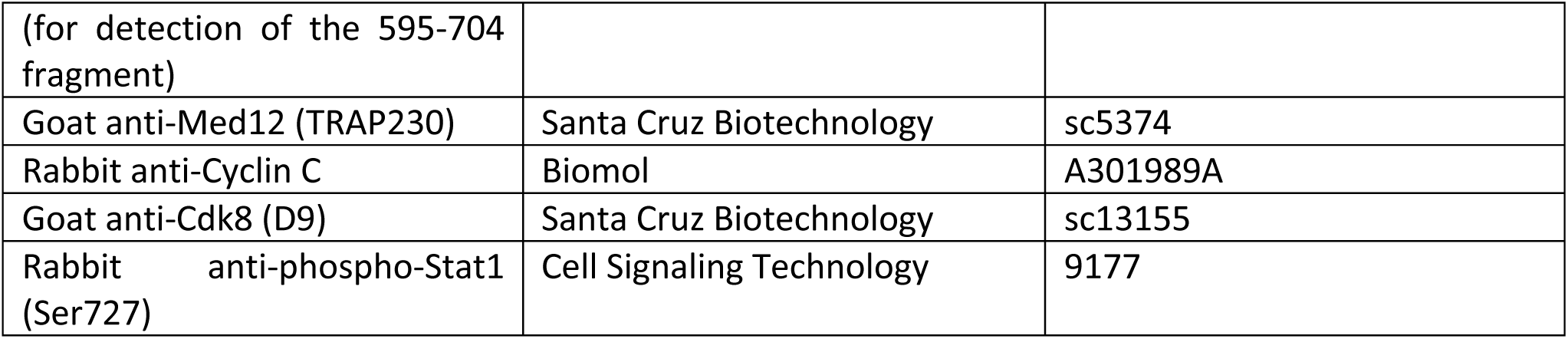

#### Commercial proteins

| Substrate | Company | Ordering number |
| --- | --- | --- |
| Recombinant Human RNA polymerase II CTD repeat | Abcam | AB81834 |

#### siRNAs

| siRNA | Name | Ordering number |
| --- | --- | --- |
| MTBP-1 | MTBP si2 On-TargetPLUS | J-013953-06-0050, Dharmacon |
| MTBP-2 | MTBP si4 On-TargetPLUS | J-013953-08-0050, Dharmacon |
| Med12-1 | siGENOME Human MED12 siRNA 2 | D-009092-02-0020, Dharmacon |
| Med12-2 | siGENOME Human MED12 siRNA 4 | D-009092-04-0020, Dharmacon |
| Cyclin C-1 | siGENOME CCNC(human) 06 | D-003209-06-0002, Dharmacon |
| Cyclin C-2 | siGENOME CCNC(human) 21 | D-003209-21-0020, Dharmacon |
| Cdk8-1 | siGENOME Human CDK8 (1024) siRNA 1 | D-003242-05-0020, Dharmacon |
| Cdk8-2 | siGENOME Human CDK8 (1024) siRNA 3 | D-003242-07-0020, Dharmacon |
| Control siRNA | GL2 control siRNA | CTM-430537, Dharmacon |

#### Cell culture chemicals

| Name | Ref.-nr, Vendor |
| --- | --- |
| Human IFN-gamma Recombinant Protein | PHC4031, Fisher Scientific |
| MSC2530818 | HY-101611, MedChemExpress |
| CCT-251921 | HY-19984, MedChemExpress |
| Senexin A | 4875, Tocris Bioscience |
| BRD6989 | HY-122586, MedChemExpress |

## Supporting information

Suppplementary Figures and Legends

## Ackknowledgements

We thank Christian Müller, Jenny Bormann and Svenja Heimann for excellent technical assistance. The lab of DB was supported by funding of the Deutsche Forschungsgemeinschaft, by FOR 2800 (project grant BO3385/4-1), and DFG project 3385/3-1. DB, ESG, FK and MK were supported by the SFB1430 consortium (Project-ID 424228829, subprojects A05, B04, Z03, B01).

