## Supplementary material for "MTBP allosterically activates Cdk8-CycC kinase activity": Suppplementary Figures and Legends

### Figure S1

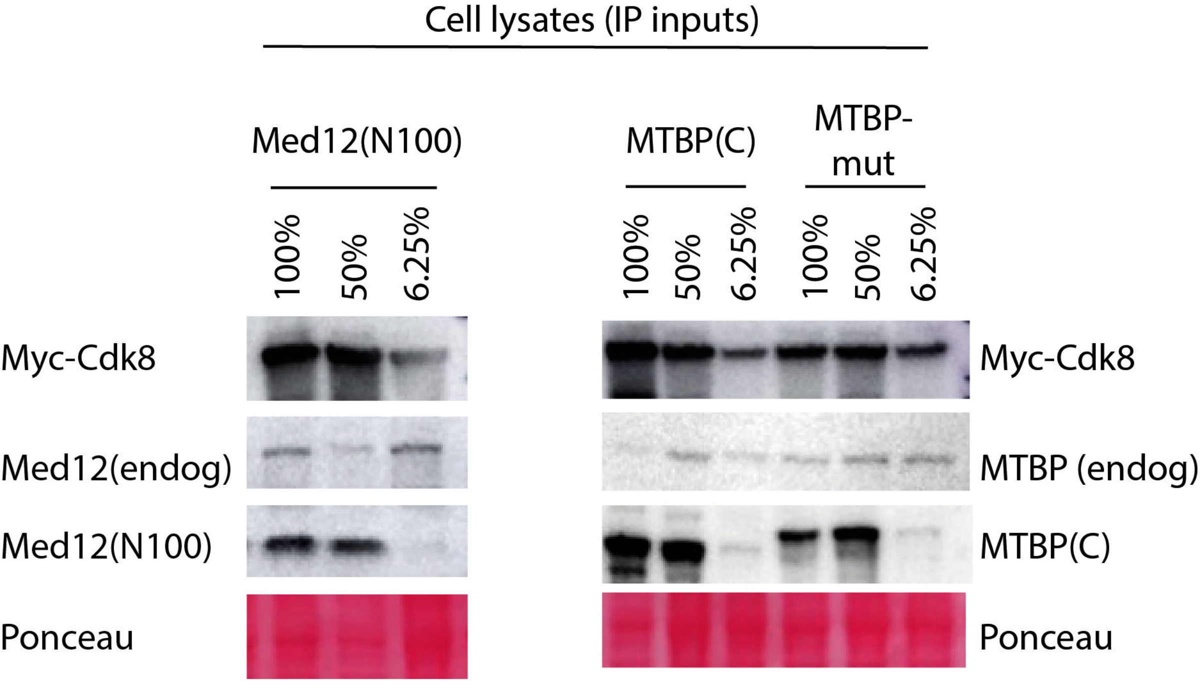

Cell lysates (IP inputs)
Med12(N100)
MTBP(C)
MTBP-mut
100%
50%
6.25%
100%
50%
6.25%
100%
50%
6.25%
Myc-Cdk8
Med12(endog)
Med12(N100)
Ponceau
Myc-Cdk8
MTBP (endog)
MTBP(C)
Ponceau

Figure S1:

Cell lysate input samples for IP experiment shown in Figure 1A

Fig. S2

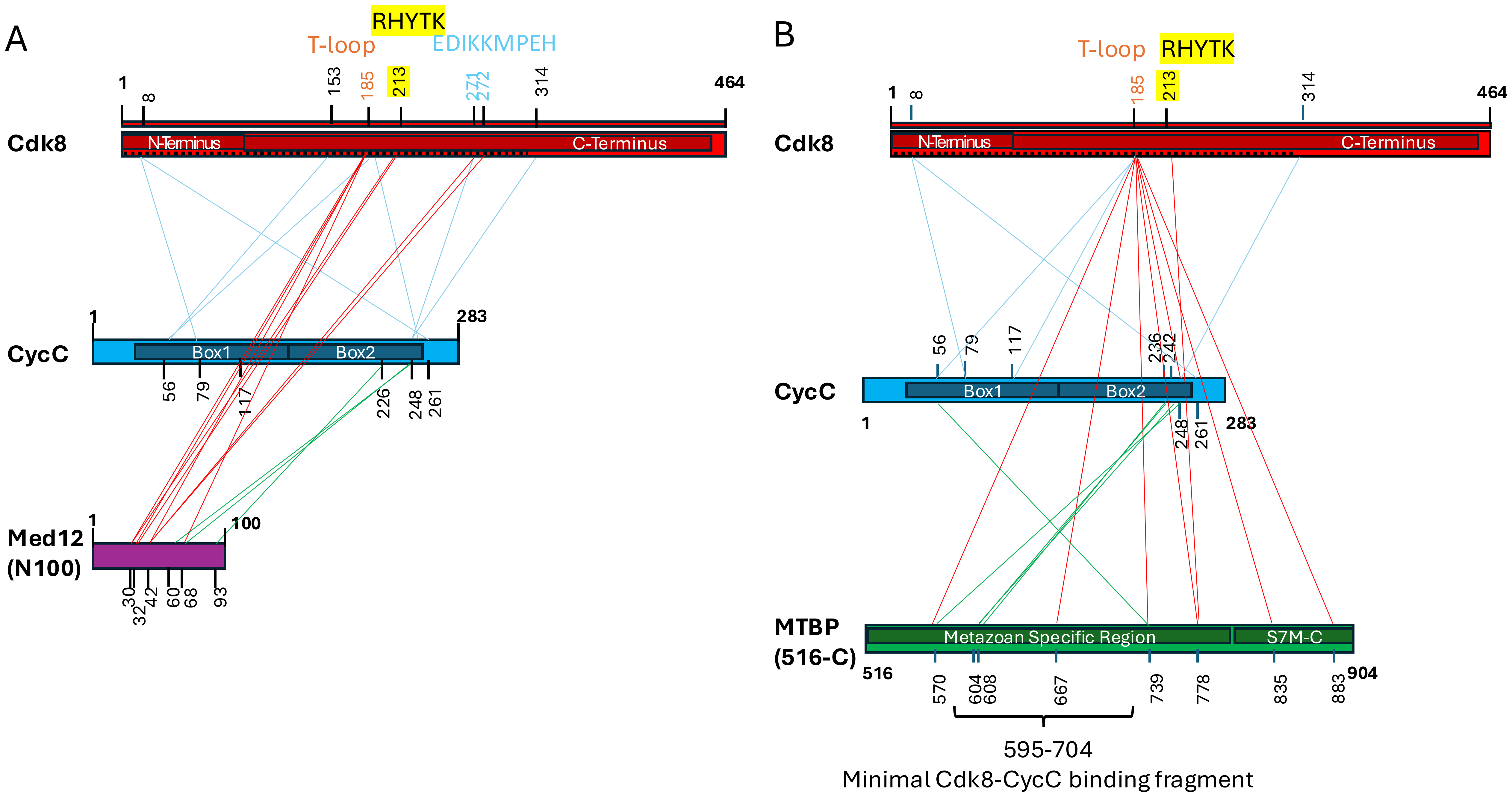
A
RHYTK
T-loop
EDIKKMPEH
1
8
153
185
213
271
272
314
464
Cdk8
N-Terminus
C-Terminus
1
283
CycC
Box1
Box2
56
79
117
226
248
261
1
100
Med12
(N100)
30
32
42
60
68
93
B
T-loop
RHYTK
1
8
185
213
314
464
Cdk8
N-Terminus
C-Terminus
56
79
117
236
242
CycC
Box1
Box2
1
248
261
283
MTBP
(516-C)
Metazoan Specific Region
S7M-C
516
570
604
608
667
739
778
835
883
904
595-704
Minimal Cdk8-CycC binding fragment

Figure S2:

BS3 crosslinking map of recombinant CycC-Cdk8 bound to Med12(N100) (A) and MTBP(516-C) (B) shown in Fig 7A (iii) and (ii), respectively. The smallest MTBP fragment mapped to bind the kinase (amino acids 595-704) is indicated in B. Whether this smallest fragment is capable to activate the Cdk8-CycC kinase activity (Figs 7, 8) remains to be tested.
